# Anti-oxidant and *in-vitro* alpha amylase inhibitory activities of leaves and barks of *Randia dumetorum*

**DOI:** 10.1101/2020.08.10.244236

**Authors:** Deepak Timalsina

## Abstract

The process of drug discovery and development in the world over a recent year has been increasingly shaped by the formulaic approaches and natural products, prioritized by popular pharmaceutical industries. In many countries like Nepal, *Randia dumetorum* (Maidal) is one of the popular alternatives for overcoming various symptoms such as acidity, food poisoning ulcers, diabetes and ulcers. It has been using by the people because of its wide range of therapeutic uses. Though, the considerable research had been conducted to reveal the biologically active compounds and its pharmacological effect, the rich constituent of this plant and its major biological perspective was yet to be studied comprehensively. Accordingly, the main focus of this study was the alpha amylase inhibition of methanolic extracts obtained from leaves and barks of *Randia dumetorum* plant. The plant extracts were obtained by dissolving dried plant with 100% methanol overnight and evaporating to the dryness. The obtained plant extract was used to study mentioned biological activities. The results obtained showed that the extract obtained from leaves and barks showed antioxidant and alpha amylase inhibition activities. This research can be helpful to discover new types of therapeutics in the future.

## Introduction

Human relationship with nature has been long and complicated from the early hood. To this day, human life is inseparable from the rich tapestry nature which offers human existence. Overwhelming evidence from historical record suggests that all aspect of human life is greatly benefitted by understanding the ever-evolving natural world. Nepal, being rich in biodiversity believed to have various medicinal plants still left to be explored. One of the famous National Parks of Nepal, “Langtang National Park” is rich in various medicinal plants lies in the Sindhupalchok district. From very ancient times people are using the plant in the form of food, shelter as well as medicine. Peoples from the upper hilly region of Sindhupalchok use this plant for the treatment of various symptoms such as food poisoning, acute stomach pain due to gastritis and ulcers. The World Health Organization (WHO) has estimated that 80% of earth inhabitants rely on traditional medicine for their primary health benefits the use of plant extract and their active components.

An ancient system of medicine adopted in India, “Ayurveda” believes in medicinal plants to treat various diseases which are considered active due to chemical constituents and are designated as “Rasayana” as one of the clinical specialties. Rasayana is a specialized procedure practiced in the form of dietary regimen promoting good habits beyond drug therapy. The propose of Rasayana is the prevention of disease and counteracting the aging process. Around 34 plants are identified as Rasayana in Ayurveda, out of which some have been specifically investigated. ^1^

*Randia dumetorum* (RD) is a large thorny shrub belonging to the family Rubiaceae. Leaves are simple, obovate, wrinkled, shiny and pubescent. Flowers are white, Fragrant, solitary, seen on at the end of short branches of these plants while fruits globose, smooth berries with longitudinal ribs, yellow when ripe. Seeds many compressed embedded in the dark fetid pulp.^2^ This plant is widely distributed in the hilly region of Nepal from 3000-4000 ft. RD was reported to contain mannitol, iridoid-10 methyloxioside, coumarin glycosides, triterpenoid glycosides, randianin and saponins named dumentoronin.^3,4^

The fruits are well-known folk medicines for their antispasmodic, anti-dysenteric and antifertility properties^1,5^. TLC analysis revealed the number of tri-terpene saponins present in the *Randia dumetorum* fruit extract. 10-methylioxide, an iridoid glucoside have been isolated from the extract of *Randia dumetorum*.^6^ Triterpene glucoside called randianin is present in the methanolic extract of *Randia dumetorum* fruit extract.^7^ The present proposed research is focused on the collection of leaves, barks, and fruits of *Randia dumetorum* from the upper region of Sindhupalchok district, Nepal to evaluate the α-amylase inhibition activities.

## Materials and methods

### Plant collection, identification, and processing

Roots, barks, and leaves of *Randia dumetorum* was collected from Sindhupalchok district of Nepal and identified at Central Department of Botany, Tribhuvan University. Collected materials was dried and shade to get powder.

### Extraction

The extraction was carried out either by cold percolation or by Soxhlet apparatus. The dried powdered sample was dipped in methanol for about 72 hours. The extract will be filtered through cotton wool and then through Whatman filter paper. The collected extract was dried by using rotary evaporator.

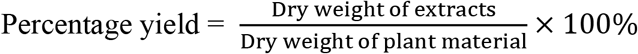

### Phytochemical screening and its analysis

The method employed for phytochemical screening and analysis of compounds present in plant extracts like Alkaloids, Flavonoids, terpenoids, and Glycosides was detected by using specific reagents as well as total phenolic and total flavonoids were estimated by following standard procedures adopted. ^8,9^

### Detection of flavonoids

Alkaline reagent test: In 5mL of methanolic extract solution 2 mL of 2 % NaOH was added, if presence of intense yellow color appears and after addition of few drops of dilute HCl turns colorless, indicates presence of flavonoid.

### Detection of terpenoids

Salkowski test: 5 mL methanolic extract was added with 2 mL of chloroform and 3 mL of concentrated sulphuric acid, appearance of reddish-brown color indicates the presence of terpenoids.

### Detection of phenols

Ferric chloride test: 50 mg extracts was taken and mixed with 5 mL of distilled water, and then few drops of 5 % ferric chloride solution was added. Presence of dark green color indicates phenol compound.

### Detection of alkaloids

50 mg extract was taken and mixed with 3 mL of diluted hydrochloric acid stirred and filtered. The filtrate was tested carefully with Mayer’s test.

Mayer’s test: Mayer’s reagent (2-3 drops) was added by the side of test tube to few mL of filtrate, and if formation of white creamy precipitate indicates presence of alkaloids.

### Detection of carbohydrates

Fehling 1 and Fehling 2 reagent was mixed in equal volume and then 2 mL of it was mixed with crude extracts. It was boiled gently and observes for brick red color at the bottom of the test tube which indicates positive test.

### Detection of glycosides

0.5 gm of extracts was taken and dissolves in 2 mL glacial acetic acid and then one drop of ferric chloride solution was added. 1 mL concentrated sulphuric acid was added to the mixture. Brown ring at the interface indicated the presence of glycosides.

### Detection of saponins

Foam test: 50 mg extract was diluted with distilled water and the volume was maintained to 20 mL. Then the suspension was shaken vigorously in a graduated cylinder for 15 minutes. A 2 cm layer of foam indicated the presence of saponins.

### Determination of total phenol content

Preparation of Reagents: 1 M sodium carbonate was prepared by dissolving 5.29 gram of sodium carbonate in 50 mL distilled water. The 1:10 v/v Folin-ciocalteu reagent was prepared by diluting 10 mL of commercially available F-C reagent in 100 mL of distilled water.

### Preparation Standard gallic acid solution

Stock solution of 500 μg/mL was prepared by dissolving 5 mg of gallic acid in 10 mL of ethanol. The stock solution was diluted to prepare final concentration of 10, 20, 30, 40, 50, 60, 70, and 80 μg/mL. The solution used for the test was freshly prepared.

### Preparation of plant extracts

The plant extracts was prepared 500 μg/mL by diluting the stock solution of 50 mg/mL in 50 % DMSO solution.

### Procedure

Total phenol content of the extracts was measured using Folin-Ciocalteu reagent by 96 well plate methods which was modified from colorimetric method. At first 20 μL of different concentration of standard 10, 20, 30, 40, 50, 60, 70 and 80 μg/mL gallic acid was loaded on 96 well plate in triplicate. Then 20 μL of plant sample of 500 μg/mL was loaded on 96 well plate in triplicate. And then in each well containing standard and sample 100 μL Folin-ciocalteu followed by 80 μL Na2CO3 was added separately. It was left in dark for 15 minute and and after 15 minute absorbance was taken at 765 nm using micro-plate reader. (Epoch2, BioTek, Instruments, Inc., USA). Gallic acid was used for constructing the standard curve. And the total polyphenolic compound concentration in the extracts was expressed as milligrams of gallic acid equivalent per gram of dry weight (mgGAE/g) of the extract using gallic acid standard curve. ^10^

### Determination of total flavonoid content

Preparation of Reagents: 10% Aluminum trichloride was prepared by dissolving 1 gram of AlCl3 in 10 mL distilled water. Similarly, 1 M potassium acetate was prepared by dissolving 0.98 grams of potassium acetate into 10 mL distilled water.

### Preparation Standard quercetin solution

Stock solution of quercetin was prepared by dissolving 1 mg of quercetin into 10 mL methanol (0.1 mg/mL). The final concentrations of the standard solution were prepared 10, 20, 30, 40, 50, 60, 70 and 80 μg/mL by diluting the stock solution.

### Preparation of plant extracts

The plant extracts were prepared 500 μg/mL by diluting the stock solution of 50 mg/mL in 50% DMSO solution.

### Procedure

Total flavonoid content of the extracts was determined by 96 well plate method which was modified from colorimetric method. First 130 μL of different concentration of standard quercetin 10, 20, 30, 40, 50, 60, 70 and 80 μg/mL was loaded on 96 well plate in triplicate. Then 20 μL of plant sample of 500 μg/mL was loaded on 96 well plate in triplicate. Distilled water of 110 μL was added in each well containing plant sample maintaining 130 μL of final volume. Then 60 μL ethanol, 5 μL AlCl3 and 5 μL potassium acetate was added in each well containing standard and plant sample, separately. It was left in dark for 30 minutes and after 30 minutes absorbance was taken at 415 nm using micro-plate reader. ^11^ (Epoch2, BioTek, Instruments, Inc., USA).

### Determination of antioxidant activity

Preparation of DPPH solution (0.1 mM) 0.1 mM DPPH solution was prepared by dissolving 3.9 mg DPPH in 100 mL methanol and covered with aluminum foil.

### Preparation of quercetin solution

Stock solution of quercetin was prepared by dissolving 2 mg of quercetin in 2 mL of methanol. Then final concentration of 20, 10, 5, 2.5 and 1.25 μg/mL were prepared by diluting the stock solution of 1 mg/mL.

### Preparation of plant extracts

Stock solution of 50 mg/mL was prepared by dissolving 50 mg of plant extracts in 1 mL of DMSO. The final concentrations of plant extracts were prepared 1000, 500, 250, 125 and 62.5 μg/mL in 50% DMSO solution.

### Procedure

Antioxidant activity of the extracts was determined by 96-well plate method which was modified from colorimetric method. For DPPH test, quercetin of 20 μg/mL was used as positive control and 50% DMSO was used as negative control. The positive control quercetin, negative control DMSO and plant samples were loaded 100 μL in 96 well plate in triplicate. Then 100 μL of DPPH reagent was added in each well, it was incubated for 30 minutes in dark. After 30 minute absorbance was taken at 517 nm using micro-plate reader.^12^

The ability to scavenge the DPPH radical was calculated by using the following equation:

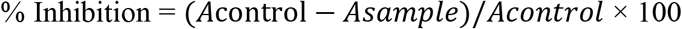

### Determination of α-amylase inhibiting properties

### Preparation of dinitrosalicylic acid

The dinitrosalicylic acid (DNSA) color reagent was prepared using 96 mM 3,5-dinitrosalicylic acid of 0.438g dissolved in deionized water (20 mL) with the help of magnetic stirrer.

### Preparation of sodium potassium tartarate

12g of sodium potassium tartarate was taken to prepare 5.31 M sodium potassium tartrate in 2 M NaOH (8 mL) and deionized water (12 mL) (Nickavar et al. 2008).

### Preparation of enzyme

1 mg/ml (9 unit) α-Amylase was prepared and finally diluted to prepare 8 unit by dissolving in 20 mM phosphate buffer (pH 6.9, containing 6.7mM NaCl).

### Preparation of plant extract

Plant extracts were dissolved in deionized water with 1 % DMSO by using vortex machine (500– 5000 ppm extracts/ 1–250 ppm for pure compounds). A 100 μL of α-amylase (8U /mL) was mixed with plant extract and incubated at 25 °C for about 30 min. A 100 μL of this mixture was mixed with starch (0.5 % w/v) solution (100μL) and incubated at 25 °C for 3 min. DNSA reagent (100 μL) was added, incubated at 85 °C for 15 min in a water bath, allowed to cool and then diluted with distilled water(900μL). Negative controls were conducted in the same manner with 1 % DMSO (100μL) in distilled water. Blanks were prepared by adding DNSA reagent prior to the addition of starch solution kept in 85 °C water bath for 15 min and then diluted with distilled water (900 μL) as before. Absorbance was measured at 540 nm. Percentage inhibition was plotted against concentration to calculate IC_50_ by software GraphPad Prism. Acarbose (1 – 100 ppm) was used as the positive control.^13,14^

## Results and discussion

### Phytochemical screening

Table showing qualitative chemical test:

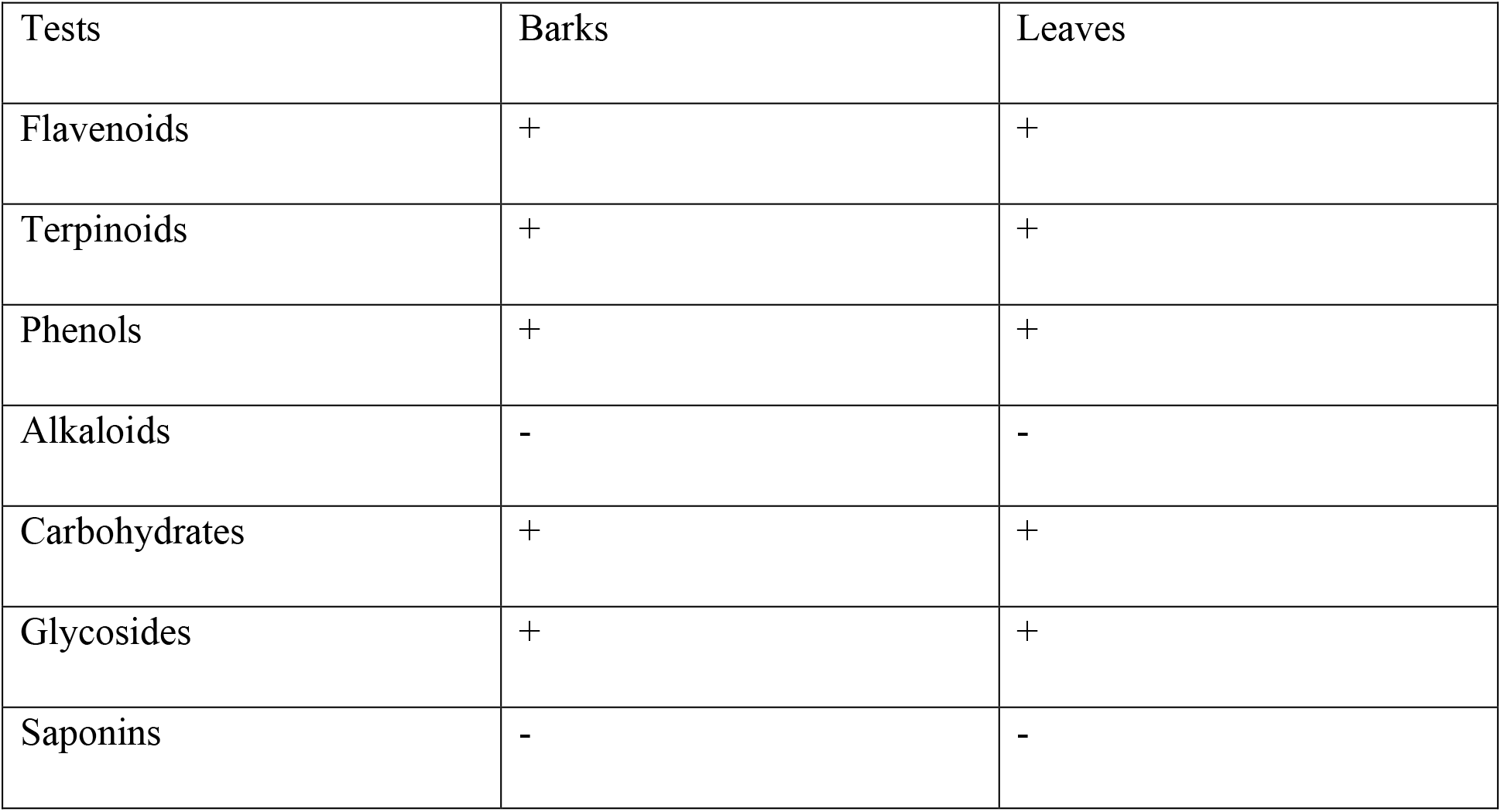

### Total phenol content

Total phenolic contain (TPC) are expressed in terms of gallic acid equivalent (mg GAE/gm dry weight of extract) with calibration curve of gallic acid (y = 0.0258x - 0.0724, R^2^ = 0.989).

| Sample | Bark | Leaves |
| --- | --- | --- |
| Total Phenolic Content | $136.66 \pm 4.38$ mg GAE/gm | $128.98 \pm 7.38$ mg GAE/gm |

### Total flavonoid content

Total flavonoid contains (TFC) are expressed in terms of quercetin equivalent (mg QE/gm dry weight of extract) with calibration curve of quercetin. (y = 0.0098x - 0.0387 R^2^ = 0.9842)

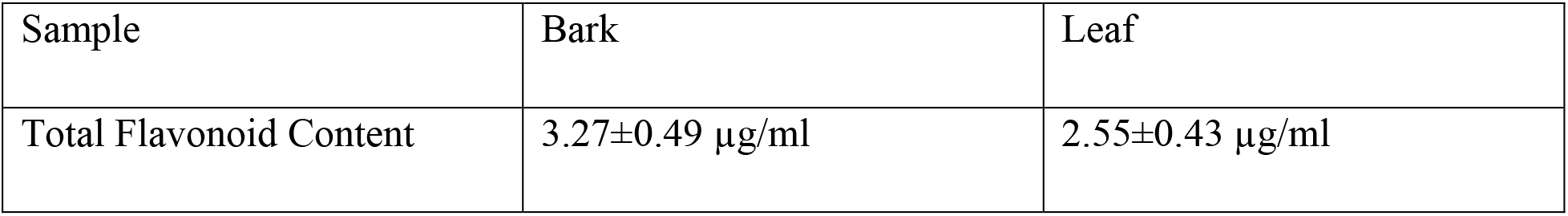

### Determination of antioxidant Capacity

#### DPPH free radical scavenging activity

1, 1-diphenyl-2-picryl hydrazyl (DPPH) was used to determine antioxidant activity of plants extracts and were expressed as IC_50_ (half inhibitory concentration). The comparative study of IC_50_ of standard and plant extract was done and tabulated as follows:

IC_50_ value of DPPH assay:

| Sample | IC<sub>50</sub> |
| --- | --- |
| Standard quercetin | 39.9±5.09 µg/ml |
| Bark | 401.6 ±4.12µg/ml |
| Leaf | 485.9±3.5µg/ml |

### α-amylase inhibitiary activities

Crude extract (500 μg/mL) of *Randia dumetorum* was evaluated for the inhibitory activity against porcine pancreatic α-amylase. The comparative study of IC_50_ of Acarabose as a positive control and plant extracts is tabulated below;

| Sample | IC<sub>50</sub> |
| --- | --- |
| Acarabose | 51.14±3.16 µg/ml |
| Bark | 203.3±4.2 µg/ml |
| Leaf | 391.5±7.1 µg/ml |

### Conclusion

The result of the present study showed that the extract of Randia dumetrum leaves and bark extracts which contains of polyphenols and flavonoids. These phytochemicals exhibited a greatest antioxidant and α-amylase inhibitory activities which participates in diabetics ageing etc. This work has gathered experimental evidence on the natural antioxidant and anti-diabetic found on *Randia dumetorum* barks and leaves. Comparatively, bark shows more active biological activities such as antioxidant and α-amylase inhibition than the leaves as total polyphenols present were greater in bark than in leaves.

## References

(1) Patil, M. J.; Bafna, A. R.; Bodas, K.; Shafi, S. In Vitro Antioxidant Activity of Fruits of Randia Dumetorum Lamk. 2014.

(2) Patel Ritesh, G.; Pathak Nimish, L.; Rathod Jaimik, D.; LD, P.; Bhatt Nayna, M. Phytopharmacological Properties of Randia Dumetorum as a Potential Medicinal Tree: An Overview. J. Appl. Pharm. Sci. 2011, 1 (10), 24–26.

(3) Kandimalla, R.; Kalita, S.; Saikia, B.; Choudhury, B.; Singh, Y. P.; Kalita, K.; Dash, S.; Kotoky, J. Antioxidant and Hepatoprotective Potentiality of Randia Dumetorum Lam. Leaf and Bark via Inhibition of Oxidative Stress and Inflammatory Cytokines. Front. Pharmacol. 2016, 7. https://doi.org/10.3389/fphar.2016.00205.

(4) Dubois, M.-A.; Benze, S.; Wagner, H. New Biologically Active Triterpene-Saponins from Randia Dumetorum. Planta Med. 1990, 56 (05), 451–455.

(5) Dubois, M.-A.; Benze, S.; Wagner, H. New Biologically Active Triterpene-Saponins from Randia Dumetorum. Planta Med. 1990, 56 (05), 451–455.

(6) Sati, O. P.; Chaukiyal, D. C.; Nishi, M.; Miyahara, K.; Kawasaki, T. An Iridoid from Randia Dumetorum. Phytochemistry 1986, 25 (11), 2658–2660.

(7) Sotheeswaran, S.; Bokel, M.; Kraus, W. A Hemolytic Saponin, Randianin, from Randia Dumetorum. Phytochemistry 1989, 28 (5), 1544–1546.

(8) Verma, R. K.; Verma, S. K. Phytochemical and Termiticidal Study of Lantana Camara Var. Aculeata Leaves. Fitoterapia 2006, 77 (6), 466–468.

(9) Tamilselvi, N.; Krishnamoorthy, P.; Dhamotharan, R.; Arumugam, P.; Sagadevan, E. Analysis of Total Phenols, Total Tannins and Screening of Phytocomponents in Indigofera Aspalathoides (Shivanar Vembu) Vahl EX DC. J. Chem. Pharm. Res. 2012, 4 (6), 3259–3262.

(10) Slinkard, K.; Singleton, V. L. Total Phenol Analysis: Automation and Comparison with Manual Methods. Am. J. Enol. Vitic. 1977, 28 (1), 49–55.

(11) Marinova, D.; Ribarova, F.; Atanassova, M. Total Phenolics and Total Flavonoids in Bulgarian Fruits and Vegetables. J. Univ. Chem. Technol. Metall. 2005, 40 (3), 255–260.

(12) Sutharsingh, R.; Kavimani, S.; Jayakar, B.; Uvarani, M.; Thangathirupathi, A. Quantitave Phytochemical Estimation and Antioxidant Studies on Aerial Parts of Naravelia Zeylanica Dc. Int. J. Pharm. Stud. Res. 2011, 2 (2), 52–56.

(13) Wickramaratne, M. N.; Punchihewa, J. C.; Wickramaratne, D. B. M. In-Vitro Alpha Amylase Inhibitory Activity of the Leaf Extracts of Adenanthera Pavonina. BMC Complement. Altern. Med. 2016, 16 (1), 466.

(14) Senger, M. R.; Gomes, L. da C. A.; Ferreira, S. B.; Kaiser, C. R.; Ferreira, V. F.; Silva Jr, F. P. Kinetics Studies on the Inhibition Mechanism of Pancreatic α-Amylase by Glycoconjugated 1H-1, 2, 3-Triazoles: A New Class of Inhibitors with Hypoglycemiant Activity. ChemBioChem 2012, 13 (11), 1584–1593.

